# Source tracing of *Leishmania donovani* in emerging foci of visceral leishmaniasis in West Nepal

**DOI:** 10.1101/2023.08.22.554278

**Authors:** Pieter Monsieurs, Kristien Cloots, Surendra Uranw, Megha Raj Banjara, Prakash Ghimire, Sakib Burza, Epco Hasker, Jean-Claude Dujardin, Malgorzata Anna Domagalska

**Affiliations:** Institute of Tropical Medicine, Antwerp, Belgium; BP Koirala Institute of Health Sciences, Dharan, Nepal; Central Department of Microbiology, Tribhuvan University, Kathmandu, Nepal; London School of Hygiene and Tropical Medicine, United Kingdom

## Abstract

We sequenced *Leishmania donovani* genomes in blood samples collected in emerging foci of Visceral Leishmaniasis, in West Nepal. We detected lineages which are very different from pre-elimination main parasite population: a new lineage and a rare one previously reported once in East Nepal. We highlight the need for genomic surveillance.

---

For decades, the Indian sub-continent (ISC) was the most endemic region for Visceral Leishmaniasis (VL) in the World. In 2005, a regional elimination program was launched in India, Nepal and Bangladesh, aiming to reduce VL anual incidence to less than one case per 10 000 at (sub-)district levels (1). Before the start of the elimination program, VL in Nepal was mainly confined to 12 VL endemic districts (out of a current total of 77 districts), located in the lowlands in the east of the country. Recently, VL cases have spread from east to west in the country, as well as from lowlands to hilly and even mountainous areas, resulting in a current total of 23 official VL endemic districts, with many more districts reporting likely indigenous cases (1). Cutaneous leishmaniasis (CL) is also becoming more common (2) and even combined cases of CL and VL are reported, without any information to date on the parasite species and genotype involved. There is clearly a need for a post-elimination surveillance system adapted to this new epidemiological profile.

Molecular surveillance of infectious diseases may provide information of highest relevance for control programs, such as (i) following the evolution of epidemics in time and space, (ii) characterization of new transmission cycles, (iii) outbreak studies and source identification and (iv) detection of new variants with new clinical features (3). Currently, no molecular surveillance is being implemented for leishmaniasis in the world, despite the existence of suitable technologies. We previously showed the feasibility and significant added value of direct whole genome sequencing (SureSelect-sequencing, SuSL) of *L. donovani* in host tissues, without the need for parasite isolation and cultivation (4).

Here, we demonstrate the proof-of-principle of SuSL for genome surveillance of leishmaniasis, in the context of the reported expansion of VL to the West of Nepal. Blood samples were collected in 2019 and stored on DNA/RNA Shield (Appendix). Three samples with the highest amounts of DNA, positive for *Leishmania* and originating from 3 different districts (Dolpa, Darchula and Bardiya, Table S1 and Fig. S1, Appendix) were sequenced, with a high genome coverage (Appendix) and compared to our database of *L. donovani* genome sequences in the ISC. The latter originated from 204 cultivated isolates (2002-2011) from Nepal, India and Bangladesh (5), 52 clinical samples (2000-2015) from Nepal (4) and 3 isolates (2002, 2010) from Sri Lanka (6,7). Altogether, these earlier studies reported 4 main genotypes: (i) a large ‘core’ group (CG), genomically very homogeneous, in the lowlands of India, Nepal and Bangladesh, (ii) a small ‘ISC1’ population, genomically very different from CG, in hilly districts of Eastern Nepal, (iii) a single divergent Nepalese isolate, BPK512 and (iv) a Sri Lanka (SL) cluster. Our new phylogenomic analyses (Fig.1) reveal that the samples from the three new foci from Western Nepal are clearly distinct from CG and SL: one ISC1-related lineage (024) had never been reported previously, while the two other lineages (022 and 023) clustered together with BPK512.

**Fig.1.**
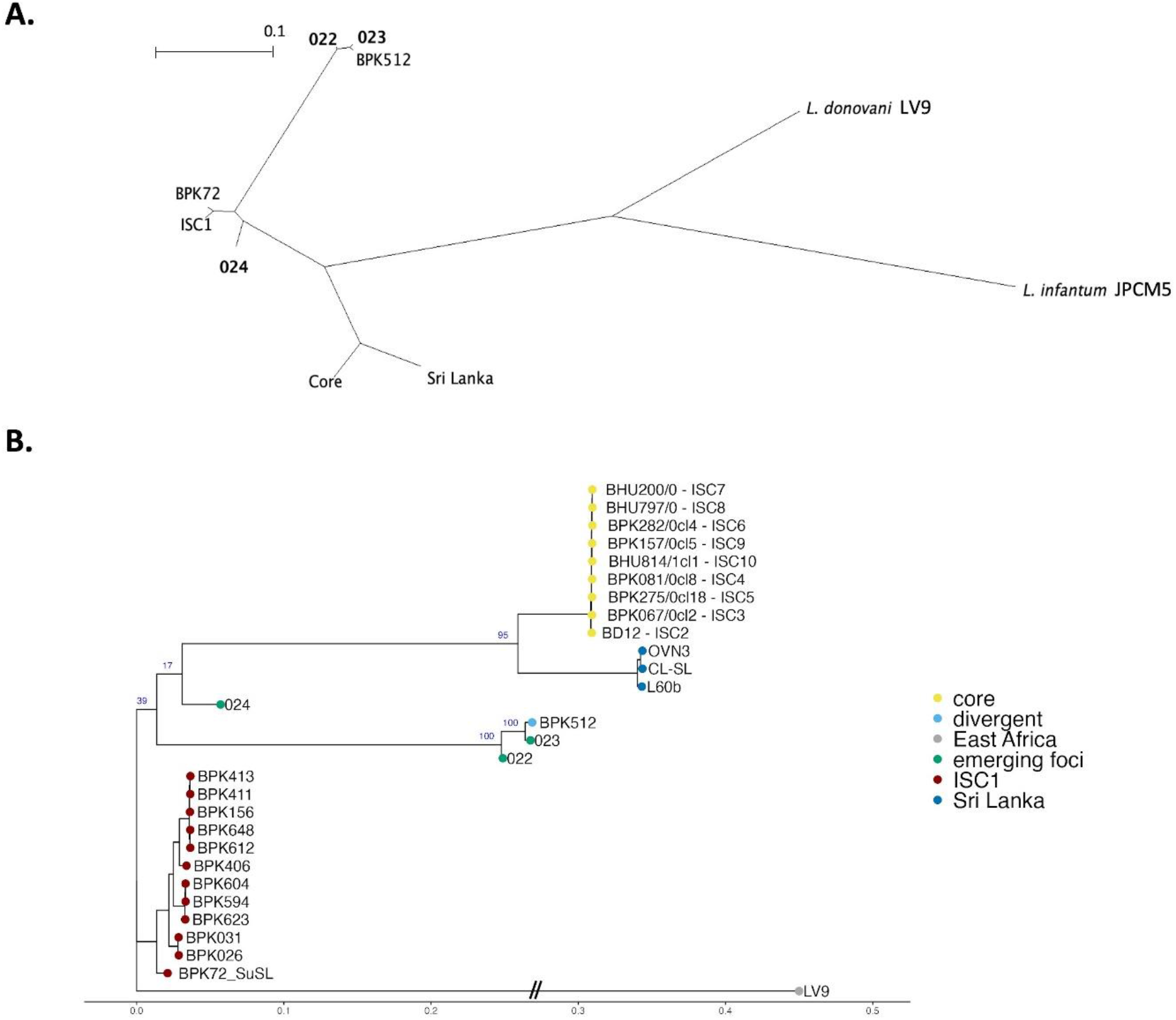
Phylogenetic analysis based on genome-wide SNPs using RAxML. **A**. Unrooted phylogenetic network of the *L. donovani* complex. Samples representing the emerging foci are indicated in bold. **B**. Rooted phylogenetic tree of reference strains of *L. donovani* from the Indian Sub-continent, showing the branching of three samples (022, 023, and 024) originating from emerging foci. Important bootstrap values are indicated in blue on the branches. The West-African LV9 strain is included as an outgroup. BPK72_SuSL represents an ISC1 sample analyzed using SureSelect, confirming that the branching of the emerging foci is not a result of a technical artifact.

It is too early to conclude that ISC1-related (024) and BPK512-like (022, 023) parasites are expanding, spreading and replacing CG in a post-elimination phase. However, a study based on single locus genotyping showed a much higher proportion of ISC1 and unclassified genotypes (and a strong decrease of CG) in 2012-2014 in comparison to 2002-2011 (8). Considering the genomic differences between these lineages and CG, we recommend a particular attention to the further evolution of parasites in the ISC. Our previous work evidenced several important functional differences between isolates from ISC1 and CG (Annex) and we found here missense mutations in 8/10 genes previously shown to be involved in *L. donovani* drug resistance (Annex). Interestingly, these genetic variants are common in the ISC1 group and in the BPK512, but never found in CG parasites. Without experimental confirmation, it is difficult to speculate about the exact impact of these mutations on the resistance to antileishmanial drugs, but it is clear that these parasite are genetically (and likely functionally) very diverse from the CG parasites, which were the main target of the recent elimination efforts.

Molecular surveillance requires a method applicable on routine samples collected in any type of field settings. Here, we demonstrate that small amounts of blood from routine examination of VL patients taken in treatment centers could be successfully used for direct, sensitive and untargeted whole genome analysis of *Leishmania*. Our optimized SuSL protocol allows highly discriminatory genotyping and thanks to its high genome coverage, it allows targeted analysis of the genetic variation within selected loci as well as untargeted searching for new markers related to a clinical or epidemiological question. Altogether, our results support the need for genomic surveillance of VL, in particular in the context of the current elimination program in the Indian subcontinent and demonstrate the applicability of SuSL to molecular surveillance of blood.

## Supporting information

annex

## Acknowledgements

Genomic sequence reads of the parasites from the three new foci are available on the European Nucleotide Archive (https://www.ebi.ac.uk/ena) under accession no.

PRJNA991731. This study was financially supported by the Belgian Directorate-General for Development Cooperation (program FA4), and the UK Department for International Development (KALACORE project).

## About the author

Dr Pieter Monsieurs is senior scientist in computational biology at ITM, expert in genomics and transcriptomics of microbial organisms. He contributed to several studies on the genomic diversity of *Leishmania, Trypanosoma and Plasmodium*, in molecular epidemiology, evolutionary and experimental contexts.

