## Supplementary material for "Source tracing of *Leishmania donovani* in emerging foci of visceral leishmaniasis in West Nepal": annex

**Appendix**

This document contains:

A1. Clinical procedures and data, including summary of clinical and epidemiological data from patients (Table S1) and a map showing their geographical origin (Fig.S1)

A2. Laboratory procedures

A3. Bioinformatic procedures

A4. Results from the analyses of specific genes reported to be involved in drug resistance (Fig.S2)

A5. Functional differences previously reported between ISC1 and CG isolates

A6. References

### A1. Clinical procedures and data

Ethical approval was obtained from the Institutional Review Committee of B.P. Koirala Institute of Health Sciences, Dharan, Nepal, as well as from the Nepal Health Research Council, Kathmandu, Nepal. In addition, ethical approval was obtained by the Institutional Review Board of the Institute of Tropical Medicine, Antwerp, and the Ethics Committee of the University Hospital of Antwerp, Belgium. Collaborating VL treatment centers were asked to collect a 2ml venous blood sample from all consenting newly diagnosed VL patients prior to the start of any treatment in January and February 2019. VL was diagnosed in line with the clinical algorithm recommended in the National Guidelines, i.e. fever > 2 weeks in combination with splenomegaly and a positive rK39 Rapid Diagnostic Test (RDT). In addition, information was collected on basic demographic factors as well as the village of residence and travel history in the two years prior to the start of symptoms. Samples were collected in DNA/RNA shield and stored at room temperature until transportation to the laboratory facilities of the Central Department of Microbiology at Tribhuvan University, Kathmandu, Nepal, where they were stored at -20°C until DNA extraction. DNA extracts were shipped to the laboratory facilities of the Institute of Tropical Medicine for further sequencing.

**Table S1.** Clinical and epidemiological data of patients; VL, Visceral Leishmaniasis; PKDL, Post Kala Azar Dermal Leishmaniasis; NA, not applicable; L-AmB, Liposomal Amphotericin B; PMM, Paromomycin

| **Sample code** | **022** | **023** | **024** |
|---|---|---|---|
| **Sequencing code** | 105328-001-022 | 105328-001-023 | 105328-001-024 |
| **date_sample_collection** | 21/01/2019 | 24/01/2019 | 07/02/2019 |
| **District** | Dolpa | Darchula | Bardiya |
| **Village** | Se-Phoxundo rural Municipality | Juga Rural Municipality | Madhuwan Municipality |
| **type_disease** | VL | VL | VL |
| **Past history of VL or PKDL** | No | Past history of VL x 6 Month | No |
| **past_drug_used** | NA | L-AmB | NA |
| **current_drug_used** | L-AmB | L-AmB + PMM | L-AmB |
| **date_treatment_start** | 21/01/2019 | 24/01/2019 | 07/02/2019 |
| **initial_outcome** | improved | improved | improved |
| **final_outcome** | no data | cured | cured |
| **travel_history_VL endemic areas** | travelled history to Surkhet districts | no travel history to VL endemic areas in Nepal & India | History of travel to Uttrakhand, India |

**Fig. S1.** Geographical origin of the three 2019 samples. The map shows the 77 districts of Nepal and those in which parasites of the core group (vertical hatched) and ISC1 (dotted) were detected from 2000 to 2015. Map was done with qGIS version 3.28.4.

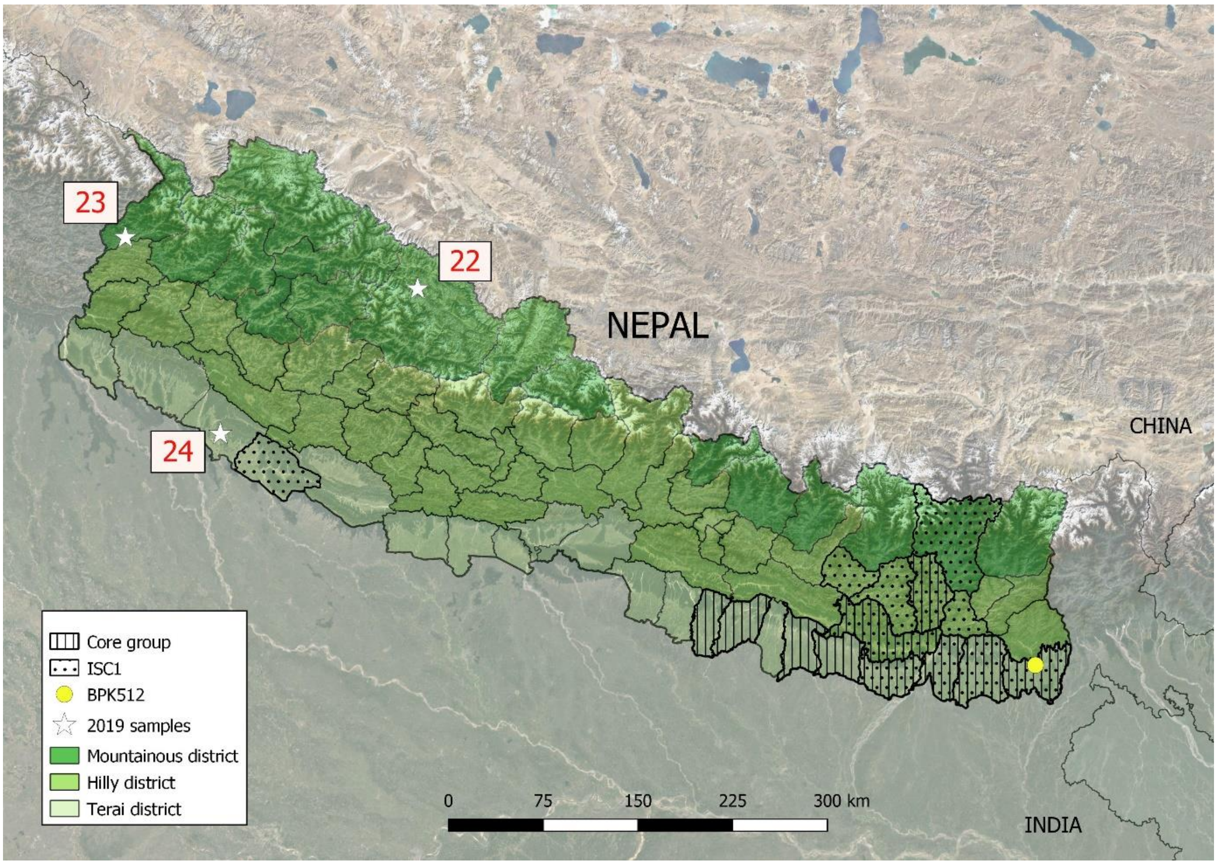

### A2. Laboratory procedures

#### DNA extraction

200µl of blood mixed with 200µl DNA/RNA Shield was used to extract DNA with QIAamp DNA Mini Kit (Qiagen) according to manufacturer's instructions with the following modifications: 30µl Proteinase K and 300µl ethanol were used instead of 20µl and 200µl, respectively. DNA concentration was verified using the Qubit broad-range DNA quantification kit (Thermo Fisher Scientific), and the % of Leishmania DNA in the samples was estimated using qPCR as described previously (1). The Leishmania % in the selected samples was 0.13, 0.16 and 0.03 in samples 022, 023, and 024, respectively. SureSelect (Agilent Technologies) was used to capture *Leishmania* genomic DNA following standard SureSelect XTHS Target Enrichment system protocol for Illumina Multiplexed Sequencing platforms. Prior the genome capture, DNA was concentrated using AMPure XP beads (Beckman, Coulter) to obtain approximately 10ng of total genomic DNA in 7 µl that was subjected to enzymatic fragmentation using SureSelect XT HS Fragmentation kit (Agilent Technologies). Custom designed oligonucleotide baits were used at 1:10 stock dilution. Sequencing was conducted on the Illumina NovaSeq platform using 2x150 bp paired reads at GenomeScan (Netherlands), for which 51.49, 51.59 and 50.35 million raw reads were obtained for sample 022, 023 and 024 respectively.

**A3. Bioinformatic procedures**

In addition to the newly sequenced data, additional sequencing data were obtained from previous publications: i) genomes describing the population structure of *L. donovani* in Nepal, India and Bangladesh (2), ii) sequencing data obtained from samples from Nepal using the SureSelect technology (1), similar to the approach used for the three outbreak samples in this report, iii) three genomes originating from Sri Lanka (3,4), and iv) the *L. infantum* sequencing data submitted under the accession number ERR1913337 (5). All publicly available sequencing data were downloaded using the SRAtoolkit software.

The reads were mapped to the reference genome *L. donovani* available at NCBI (accession number GCF_000227135.1) using BWA (version 0.7.17(6)) with a seed length set to 50 17 (6). Only properly paired reads with a mapping quality higher than 30 were selected using SAMtools (7). Duplicate reads were removed using the RemoveDuplicates command in the Picard software (version 2.22.4, http://broadinstitute.github.io/picard/ ). SNP calling was performed using the Genome Analysis ToolKit (GATK) (8)pipeline (version 4.1.4.1) following the GATK best practices approach: 1) GATK HaplotypeCaller enabling the production of GVCF formatted files, 2) GVCF files of all samples were combined using the GATK CombineGVCF command, 3) genotyping was performed via the GATK GenotypeGVCF command, and 4) filtering of the SNPs and indels was carried out following the "best practices" approach as suggested on the GATK support site using the SelectVariants and VariantFiltration commands. Regions in the vcf-file corresponding to known drug resistance markers were selected using BCFtools, and visualized using the pheatmap function in R .

Phylogenetic trees were constructed using RAxML (9) . First, the VCF files containing biallelic SNPs were selected using BCFtools (7) and were converted to Phylip format using the vcf2phylip.py script (https://github.com/edgardomortiz/vcf2phylip). RAxML was then executed with the GTR+G substitution model, utilizing 1000 bootstrap replicates. The *L. infantum* JPCM5 or the *L. donovani* LV9 genome was employed as an outgroup. The resulting phylogenetic trees were visualized using ggtree (10) for rooted phylogenetic trees and SplitsTree (11) for unrooted phylogenetic networks.

**A4. Results from the analyses of specific genes reported to be involved in drug resistance**

We selected 10 loci that were previously shown to be involved in *L. donovani* resistance to known antileishmanial drugs:

- Antimony: Aquaglyceroporin 1, AQP1 LDBPK_310030 (12); ABC transporter MRPA, MRPA LDBPK_230290 (13)
- Amphotericin B: sterol C5-desaturase, C5D LDBPK_231560 (14); sterol C24-methyltransferase, SMT LDBPK_362520 (14)
- Miltefosine: *Leishmania donovani* miltefosine transporter, LdMT LDBPK_131590 (15); Beta-subunit of LdMT, LdRos3 LdBPK_320540 (15) and genes part of miltefosine sensitivity locus (16): 3,2-trans-enoyl-CoA isomerase 1 and 2, TECI1 LdBPK_312320 and TECI2 LdBPK_312400; helicase-like protein, HELI LdBPK_312390; 3'-nucleotidase/nuclease, NUC LdBPK_312380.

For each of the three new *L. donovani* genomes, the sequence of the 10 loci was studied in details. The 10 loci were well-covered and in 8 out of the 10 genes we found at least one homozygous single nucleotide polymorphism (SNP), which results in a missense mutation or a frameshift in at least 2 out of 3 samples from new emerging loci (Fig.S2). No significant changes were observed for LdRos3 and SMT.

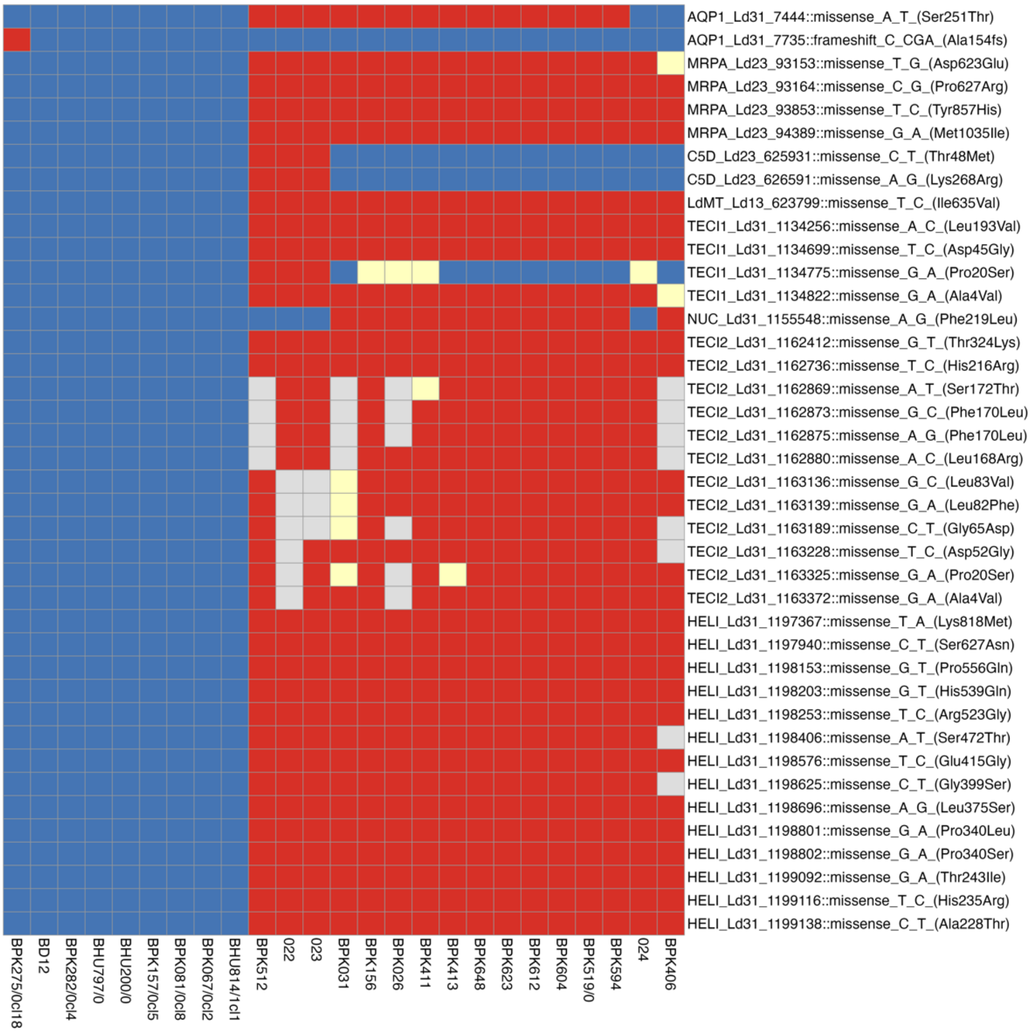


**Fig. S2:** Heatmap showing the distribution of single nucleotide polymorphisms (SNPs) in genes reported to be associated with drug-resistant phenotypes. The color scheme represents different

SNP categories: blue indicates the absence of SNPs, orange indicates heterozygous SNPs, and red indicates homozygous SNPs. The naming convention for SNPs follows the format of the gene of interest, position in the genome, type of mutation, and its effect on the corresponding protein. Samples: BPK512, CG and ISC1 isolates used for the phylogenomic analysis of Fig.1 together with the three blood samples of 2019 (022, 023, 024)

**A5. Functional differences previously reported between ISC1 and CG isolates**

Noteworthy, all results here compiled concern the analysis of isolated and cultivated parasites.

In a first study, we demonstrated that CG parasites were intrinsically more tolerant to trivalent antimonials than ISC1 ones. This phenotype was driven by the amplification of a locus containing MRPA, a gene involved in $Sb^{III}$ sequestration (13)

In a second study, we made an integrated genomic and metabolomic profiling of ISC1 vs CG isolates (17). We found several genomic differences including SNPs, CNV and small indels in genes coding for known virulence factors, immunogens and surface proteins. With respect to the metabolome, we found differences in several functional groups and pathways, essentially:

(i) **Lipid metabolism**, with 19 glycerophospholipids (GPLs) showing significantly different levels between both groups: GPLs are involved in a wide array of cellular functions including host cell infection

(ii) **Urea cycle**. In ISC1 versus CG we detected a higher concentration of citrulline and a lower concentration of argininosuccinate: Mutants for argininosuccinate synthase genes have shown a lower virulence than WT parasites.

(iii) **Nucleotide salvage pathway**. This pathway is essential, since *Leishmania* cannot synthesize the purine ring de novo and is therefore dependent on salvaging these from host purines. Our previous results suggested that ISC1 parasites might be better at salvaging nucleotides from their environment.

In a third study, we experimentally demonstrated that ISC1 and CG strains are developing similarly in natural ISC vector *Phlebotomus argentipes*, suggesting that *P. argentipes* is a fully competent vector for ISC1 parasites (18).

Altogether, these experimental studies demonstrate differences between ISC1 and CG in antimonial susceptibility and predict major functional differences, including virulence. Taking into account that ISC1 can easily be transmitted by *P. argentipes*, particular attention is required to monitor the fate of ISC1-related population in the region, especially in a post-VL elimination context.
